# The intricate triangular interaction between protective microbe, pathogen, and host genetics determines fitness of the metaorganism

**DOI:** 10.1101/2023.03.22.533850

**Authors:** Hanne Griem-Krey, Carola Petersen, Inga K. Hamerich, Hinrich Schulenburg

**Author notes:** Corresponding Author: Hinrich Schulenburg. Shared first authorship (Hanne Griem-Krey, Carola Petersen).

## Abstract

The microbiota shapes host biology in numerous ways. One example is protection against pathogens, which is likely critical for host fitness in consideration of the ubiquity of pathogens. The host itself can affect abundance of microbiota or pathogens, which has usually been characterised in separate studies. To date, however, it is unclear how the host influences the interaction with both simultaneously and how this triangular interaction determines fitness of the host-microbe assemblage, the so-called metaorganism. To address this current knowledge gap, we focused on a triangular model interaction, consisting of the nematode *Caenorhabditis elegans*, its immune-protective symbiont *Pseudomonas lurida* MYb11, and its pathogen *Bacillus thuringiensis* Bt679. We combined the two microbes with *C. elegans* mutants with altered immunity and/or microbial colonisation, and found that (i) under pathogen stress, immunocompetence has a larger influence on metaorganism fitness than colonisation with the protective microbe, (ii) in almost all cases, MYb11 still improves fitness, and (iii) disruption of p38 MAPK signalling, which contributes centrally to immunity against Bt679, completely reverses the protective effect of MYb11, which further reduces nematode survival and fitness upon infection with Bt679. Our study highlights the complex interplay between host genetics, protective microbe, and pathogen in shaping metaorganism biology.

## Introduction

The association between a host and its microbiota is central to the biology and fitness of the host shaping traits including development, physiology, immunity, reproduction, stress resistance and even behaviour [1–3]. Consequently, hosts and their associated microbes are considered as a functional unit in evolution, the metaorganism, in which the effects mediated by the microbiota are integrated into the host phenotype [4,5]. Genetic and thus phenotypic variations in the metaorganism can result from both changes in the host genome and changes in the microbiota such as acquisition of new microbes or changes in relative species abundance [2,6,7]. As a consequence, the microbiota may influence the distribution of host phenotypes within a population and thus the evolutionary trajectory of the host [6,8].

One important evolutionary advantage the microbiota provides to its host is protection against invading pathogens [9]. Microbiota-mediated protection can occur through several mechanisms, including direct interaction with the pathogen through secretion of antimicrobial products or resource competition [10–13], or indirectly through induction of the host’s immune system [14–16]. Changes in the microbiota may therefore increase host susceptibility to infections [17,18]. Moreover, the microbiota not only contains beneficial but also potentially harmful microbes. For example, *Candida albicans* is commonly found in the gut of humans but overgrowth due to changes in host immunity or microbiota cause opportunistic infections [19,20]. Additionally, pathogens have evolved strategies to promote their growth despite the microbiota or even use the microbiota to facilitate infection [21–23]. Invading pathogens may trigger changes in the microbiota causing otherwise neutral or beneficial microbes to become harmful [24,25]. Hence, beneficial and harmful microbes can interact with each other, and this interaction has consequences for the host, thereby shaping the phenotype and fitness of the metaorganism as a whole.

The host, and particularly the host immune system, directly affects both beneficial and pathogenic microorganisms. The importance of the immune system in controlling and eliminating pathogenic microbes is very well researched and forms the main basis of our current understanding of animal immunity. The host’s immune system also plays a critical role in shaping the composition of its microbiome, as demonstrated for a diversity of host systems, ranging from early branching metazoans, such as *Hydra* polyps [26], to flies and worms [27– 30], bobtail squids [31], and vertebrates and humans [32,33]. To date, it is unclear how exactly the host simultaneously influences abundance of both, the protective microbe and the pathogen, and thereby shapes metaorganism biology [34,35]. Considering the ubiquity of pathogens, this component of the triangular interaction is likely a critical factor that determines metaorganism fitness. One of the few exceptions that addressed this topic, used a laboratory-based model, consisting of the nematode host *Caenorhabditis elegans*, the protective microbe *Enterococcus faecalis*, and the pathogenic bacterium *Staphylococcus aureus* [36]. Although the two microbes are unlikely to coexist with *C. elegans* in nature, their experimental analysis yielded important insights into the principles that shape the evolution of symbiosis. Of relevance here is the finding that the host can evolve to harbour a larger number of the protective bacteria, thereby exploiting the function provided by the symbiont [37]. This effect appears to be mediated (at least to some extent) by the host immune system, for example by the lysozyme gene *lys-7*, which was identified to be part of the symbiont-dependent protective effect and which directly reduces pathogen-mediated killing and, at the same time, favours abundance of the protective symbiont over the pathogen [38]. It is currently unknown whether this host immunity-mediated effect translates into a competitive fitness advantage.

The aim of the current study is to assess the consequences of host genetics on the interaction with protective microbes and pathogens and ultimately metaorganism fitness. To address this aim, we focused on a model triangular interaction that included *C. elegans* as a host and its naturally associated protective symbiont, *Pseudomonas lurida* MYb11, as well as its pathogen *Bacillus thuringiensis*. Over the last two decades, the nematode has been used intensively for studying invertebrate immunity, with detailed information available on defence against *B. thuringiensis*, which includes insulin-like and p38 mitogen-activated protein kinase (MAPK) signalling [39–42], but not transforming growth factor beta (TGF-*β*) signalling that is involved in immunity against other pathogens [43]. More recent research efforts have been directed to the study of *C. elegans*-microbiota interactions [44]. In nature, the nematode is associated with a species-rich microbiota [45,46], which includes immune-protective microbes, such as a variety of *Pseudomonas* species [47,48]. Moreover, innate immunity [27,28,49] and microbe-microbe interactions have been shown to influence *C. elegans* microbiota composition and function [46,50]. For the current study, we used different *C. elegans* mutants, which either affect the nematode’s immune system (e.g., p38 MAPK and TGF-*β* signalling) or microbial colonisation (i.e., through disruption of the grinder in the nematode’s pharynx). The *C. elegans* mutants and wildtype were exposed in different combinations to the immune-protective symbiont *P. lurida* MYb11 and pathogenic *B. thuringiensis* MYBT18679 (Bt679), followed by assessment of microbial colonisation rates, nematode survival, nematode population growth, and importantly, nematode competitive fitness.

## Material and Methods

### Nematode and bacterial strains

The *C. elegans* strains used in this study are listed in Table 1. Prior to all experiments, *C. elegans* strains were thawed from frozen stocks, maintained on nematode growth medium (NGM) on *Escherichia coli* strain OP50, and synchronized by bleaching following standard procedures [51].

**Table 1:** *C. elegans* wildtype, nine mutant strains and one transgenic strain used in this study.

| Strain* | Genotype | Description |
| --- | --- | --- |
| N2 | wildtype | Wildtype |
| *dbl-1* | *dbl-1(nk3)* | TGF-β defective |
| *sma-6* | *sma-6(e1482)* | TGF-β defective |
| *phm-2* | *phm-2(ad597)* | Grinder defective |
| *tnt-3* | *tnt-3(ok1011)* | Grinder defective |
| *daf-2* | *daf-2(e1370)* | Insulin-like signalling defective |
| *daf-16* | *daf-16(mgDf50)* | Insulin-like signalling defective |
| *tir-1* | *tir-1(tm3036)* | p38 MAPK defective |
| *pmk-1* | *pmk-1(km25)* | p38 MAPK defective |
| *nhr-8* | *nhr-8(ok186)* | Nuclear hormone receptor defective |
| CL2122 | *dvIs15 [(pPD30.38) unc-54(vector) + (pCL26) mtl-2::GFP]* | GFP-labelled reference strain |

\*Strain names used throughout the manuscript

*Pseudomonas lurida* MYb11 was used as a representative, protective natural microbiota member of *C. elegans* [45,47]. *Bacillus thuringiensis* MYBT18679 (Bt679), which produces pore-forming toxins, was used as a pathogen. *E. coli* OP50 and the non-pathogenic *B. thuringiensis* strain 407 Cry- (Bt407) were used as control bacteria. For each experiment, MYb11 and OP50 were freshly thawed from frozen stocks and grown on tryptic soy agar (TSA) plates at 25 °C for two days. Liquid cultures were produced from single colonies grown in tryptic soy broth in a shaking incubator overnight at 28 °C and adjusted to OD60010 in phosphate-buffered saline (PBS). *B. thuringiensis* spore stocks were prepared as previously described and frozen in aliquots at -20 °C [42,47,52]. Aliquots with spore concentrations ranging from 10^9^-10^10^ particles/ml for Bt679 and 10^3^-10^4^ particles/ml for Bt407 were freshly thawed for each experiment. All experiments were performed on peptone-free nematode growth medium (PFM) to avoid germination of *B. thuringiensis* spores.

### Bacterial colonisation assay

We assessed intestinal colonisation by MYb11 following the previously described protocol [48,53,54]. Briefly, *C. elegans* were grown until the 4th instar larval (L4) stage on PFM plates inoculated with MYb11 or OP50. L4 were transferred to 60 mm PFM plates inoculated with 500 µl MYb11 or OP50 with or without *B. thuringiensis* Bt679 (1:200 if not stated otherwise). After 24 h at 20 °C, adult worms were washed off the plates using 1.5 ml M9-buffer with 0.025 % Triton-X 100 (M9-T). After 5-times washing with 1 ml M9-T, 100 µl worm pellet was transferred to a new microtube and 100 µl 10 mM tetramisole hydrochloride were added to stop pharyngal pumping and excretion of bacteria. Worms were surface sterilized using M9-T with 2 % sodium hypochlorite and washed twice in 1 ml PBS with 0.025 % Triton-X 100 (PBS-T). Next, approximately 30 worms were transferred to a new sterile 2 ml microtube. The exact worm number was recorded for calculation of the colonisation rate per worm. The volume of the transferred worms was topped up to 400 µl with PBS-T and 10-15 1 mm zirconia beads were added. 100 µl supernatant was transferred to a new sterile microtube of which 50 µl were plated onto TSA as a control for the surface sterilization process. Worms were shredded in the remaining 300 µl PBS-T using a Bead Ruptor 96 (OMNI International, Kennesaw, Georgia, USA) for 3 min at 30 Hz. 50 µl of serially diluted, homogenized worms in PBS-T (10^−1^ to 10^−3^) were plated onto TSA. After approximately 24 h at 25 °C, colonies were counted. For Bt679 infected worms, colonies of MYb11 or OP50 and Bt679 were morphologically distinguished and counted on the same plate. For all bacteria, colony forming units (CFU) per worm were calculated.

### *B. thuringiensis* survival assay

*B. thuringiensis* survival assays were performed as previously described [42,47,48] to characterise microbiota-mediated protection against Bt679 in the different *C. elegans* mutants. Briefly, L4 larvae grown on MYb11 or OP50 on PFM plates were washed five times in sterile M9-buffer. Approximately 30 washed L4 larvae were transferred to 60 mm PFM plates inoculated with 75 µl of Bt679 spore solution mixed 1:25, 1:50, 1:100, 1:200 or 1:500 with MYb11 or OP50. Plates inoculated with the non-pathogenic Bt407 mixed with MYb11 or OP50 at the highest concentration used for pathogenic Bt679 served as controls. After 24 h at 20 °C, the number of dead and alive worms was counted. Worms were considered dead if they failed to respond to light touch. The survival rate was determined as proportion of worms alive.

### Population growth rate

As a proxy for fitness of the *C. elegans* strains when challenged with Bt679, a population growth assay was performed as previously described [45,47,48,55]. Briefly, *C. elegans* were grown on MYb11 or OP50 on PFM plates until L4 stage. Three L4 larvae were picked onto 90 mm PFM plates inoculated with 1 ml Bt679 or Bt407 spore solution mixed 1:200 with MYb11 or OP50 (worms were maintained on the same non-pathogenic bacteria as during larval development). After five days at 20 °C, the worms were washed off the plates using 5 ml M9-T and directly frozen at -20 °C until scoring. Worms on plates with low worm counts were scored directly. For scoring, frozen samples were thawed, mixed thoroughly, and the worms counted in triplicates in droplets ranging from 10-100 µl depending on worm density, followed by calculation of the offspring number per worm.

### Competitive fitness assay

We assessed the effect of the different microbiota-mediated effects on competitive fitness of the *C. elegans* mutants. Competitive fitness was determined using a population growth assay where the considered *C. elegans* mutants were always competed against a reference *C. elegans* strain, the transgenic strain CL2122, which produces a strong constitutive intestinal expression of GFP at all developmental stages [56]. To control for potential effects of GFP-labelling, competition between CL2122 and wildtype N2 was additionally tested. Three *C. elegans* CL2122 and three mutant L4 worms were picked to PFM plates inoculated with MYb11 or OP50 with or without Bt679 (1:500). The competition experiment was performed either on 60 mm PFM plates on 500 µl of Bt679 mixture or, because of high progeny numbers, on 90 mm PFM plates on 1 ml of MYb11 or OP50 in the absence of Bt679. The worms were washed off the plates with or without Bt679 after five or four days, respectively. The number of fluorescent and non-fluorescent worms was counted in triplicates in droplets of 5-20 µl depending on worm density using a fluorescent dissecting scope (Leica Microsystems GmbH, Wetzlar, Germany). Competitive fitness was determined as relative proportion of non-fluorescent mutant worms compared to the fluorescent reference strain CL2122.

### Statistical analysis

Statistical analyses and figures were done using R (Version 4.1.2) [57]. Graphs were plotted using the package ggplot2 [58] and edited in Inkscape (Version 1.1.2). The raw data can be found in Supplementary Table S1. The results of the statistical tests performed on the phenotypic raw data can be found in Supplementary Table S2. Parametric tests were used if the data met the assumptions, otherwise equivalent non-parametric tests were used. Assumptions of normality of data were confirmed with Shapiro-Wilk test and equality of variances with F-tests [59,60]. A binomial generalized linear model was used to compare differences in survival [61], followed by Tukey’s test [62]. Wilcoxon Rank-Sum tests were applied to assess differences in bacterial load and population growth [63]. One Sample t-tests were used to evaluate differences between *C. elegans* strain proportions after competition to the theoretical value of 0.5, while Two Sample t-tests served to assess variation in competitive fitness on OP50 and MYb11 [64]. False discovery rate (FDR) correction was used to account for multiple testing where appropriate [65]. Observer bias was minimized by coding of samples, the compared treatments were assayed in parallel in randomized arrangements. In the initial analysis of the nine *C. elegans* mutants, one replicate was excluded from the data, because it clearly displayed an experimental error (absence of worms).

## Results

### Interactions between diverse host genes and the protective *P. lurida* MYb11 shape *C. elegans* pathogen resistance

As a first step of our study, we explored variation among all considered *C. elegans* mutants in two traits: colonisation by the protective *P. lurida* MYb11 and survival upon Bt679 infection. The results of this survey were then used to choose a subset of mutants for further analysis. The considered mutants either showed impaired immunity or impaired microbial uptake which potentially influences microbiota abundance [27,28,49,66]. We found that MYb11 colonisation was significantly higher in the mutants *dbl-1, sma-6, phm-2, tnt-3*, and *tir-1* compared to the wildtype N2 (*p* < 0.05). In contrast, *daf-2* mutants showed lower MYb11 colonisation (*p* = 0.022), while the mutants *pmk-1, daf-16 and nhr-8* did not differ from N2 (Supplementary Figure S1A, Supplementary Table S2). We further found that survival upon Bt679 infection was significantly lower in the MYb11-fed *phm-2, tnt-3, tir-1* and *pmk-1* mutants compared to N2 (*p* < 0.001), while there was no difference in survival between the mutants *dbl-1, sma-6, daf-2, nhr-8* and N2 (Supplementary Figure S1B, Supplementary Table S2). Based on these results, we selected four strains with contrasting patterns for a more in-depth characterisation, including the mutants *dbl-1* (high colonisation, high survival), *phm-2* (high colonisation, low survival), *pmk-1* (low colonisation, low survival), and the wildtype strain N2 (low colonisation, high survival).

### Infection with *B. thuringiensis* alters intestinal colonisation rates in *C. elegans*

In a bacterial colonisation assay (Figure 1A), we subsequently tested whether Bt679 infection alters the intestinal colonisation rate with MYb11 or *E. coli* OP50. On Bt679, *dbl-1* mutants showed decreased MYb11 colonisation (*p* = 0.039), and colonisation with OP50 was decreased in *phm-2* mutants (*p* = 0.039) (Figure 1B, Supplementary Table S2). For *pmk-1* mutants an opposite trend was observed, namely that MYb11 colonisation was increased on Bt679 (*p* = 0.057). Except for the *pmk-1* mutant, MYb11 colonisation was higher than OP50 colonisation in all worm strains (*p* < 0.05). For the *pmk-1* mutant, a trend towards higher MYb11 colonisation compared to OP50 colonisation was observed without Bt679 (*p* = 0.057). Albeit not statistically significant, Bt679 was found in low numbers but more in *dbl-1* and *phm-2* mutants fed with MYb11 (Figure 1C, Supplementary Table S2).

**Figure 1:**
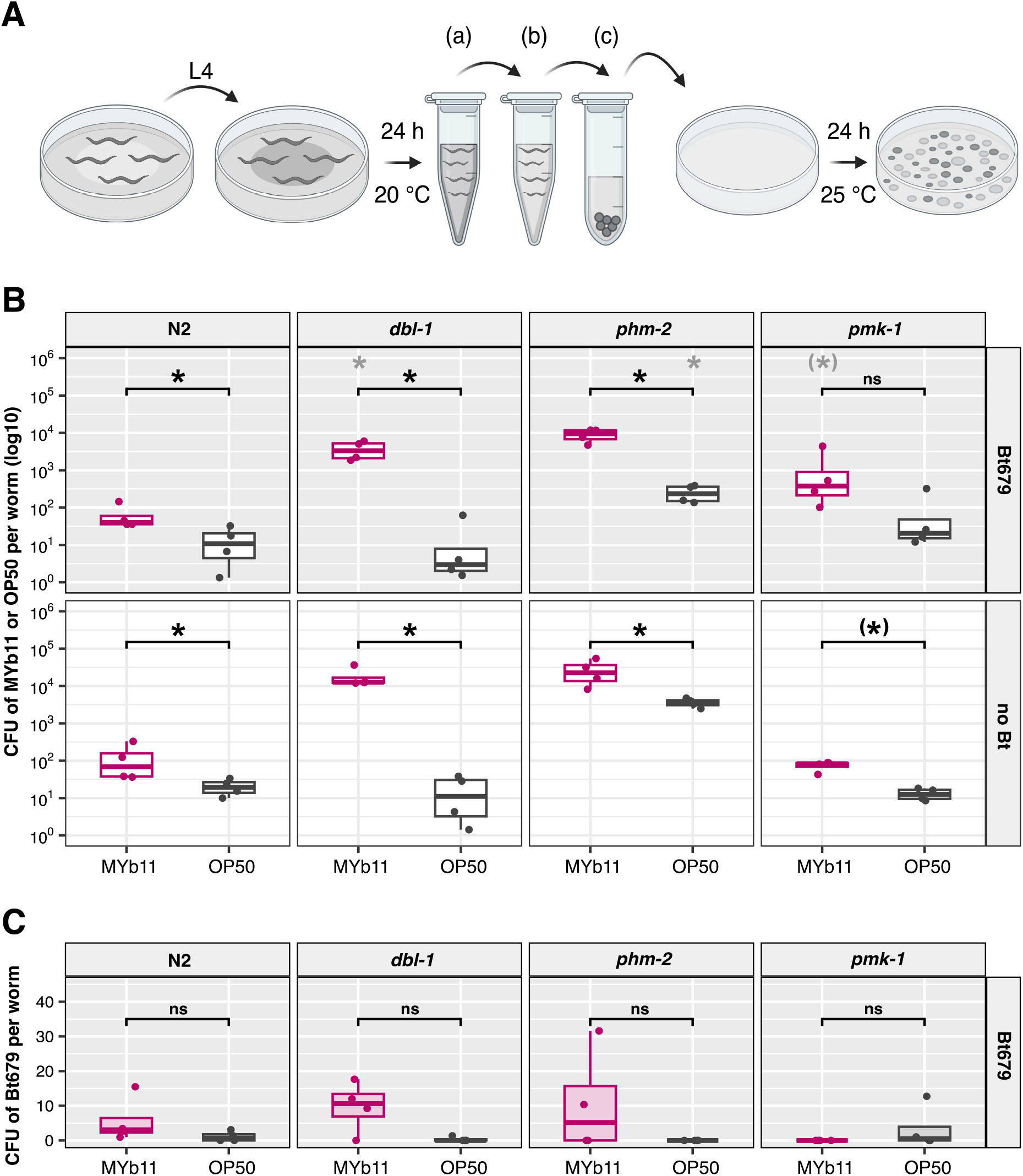
Intestinal colonisation of *C. elegans* is influenced by host genotype and *B. thuringiensis* Bt679 infection. **(A)** Synchronized *C. elegans* were grown on *P. lurida* MYb11 or *E. coli* OP50 until the L4 stage. L4 worms were transferred to the same bacteria with or without the addition of Bt679 spores (1:200). A lower Bt concentration of 1:2000 was used for the *pmk-1* mutant to account for its high susceptibility to Bt679 infection. After 24 h, adult *C. elegans* were washed (a), surface sterilized (b), and broken up (c) to release the intestinal bacteria, which were plated out to count the colony forming units (CFU). **(B)** CFU of MYb11 (pink) or OP50 (grey) per worm with Bt679 (grey panel background) or without (white panel background). Asterisks denote significant differences between MYb11 and OP50 (black) or between treatments with and without Bt679 (grey). Note that the y-axis is log10-transformed. **(C)** CFU of Bt679 per worm on MYb11 (pink) or OP50 (grey). **(B, C)** Shown are boxplots with the median as a thick horizontal line, the interquartile range as box, the whiskers as vertical lines, and each replicate depicted by a dot. Wilcoxon Rank-Sum test, corrected for multiple comparisons with FDR, ^(^*^)^*p < 0*.*1*,\**p* < 0.05, n = 4.

### Disruption of p38 MAPK pathway and grinder abrogates the protective phenotype conferred by *P. lurida* MYb11

To assess the relative contributions of host genetics and protective microbiota to pathogen resistance, we repeated the survival assay including worms grown on OP50. After Bt679 infection, survival of *C. elegans* N2 and the *dbl-1* mutant was higher on MYb11 than on OP50 (*p* < 0.001), with higher survival rates of *dbl-1* mutants compared to N2 on OP50 (*p* < 0.001) and MYb11 (*p* = 0.016) (Figure 2, Supplementary Table S2). *phm-2* mutants showed no difference in survival on MYb11 and OP50, but an overall higher survival than N2 on OP50 (*p* = 0.012). Survival of *pmk-1* mutants was generally lower than that of N2 (*p* < 0.001), with survival rates on MYb11 being significantly lower than those on OP50 (*p* < 0.001).

**Figure 2:**
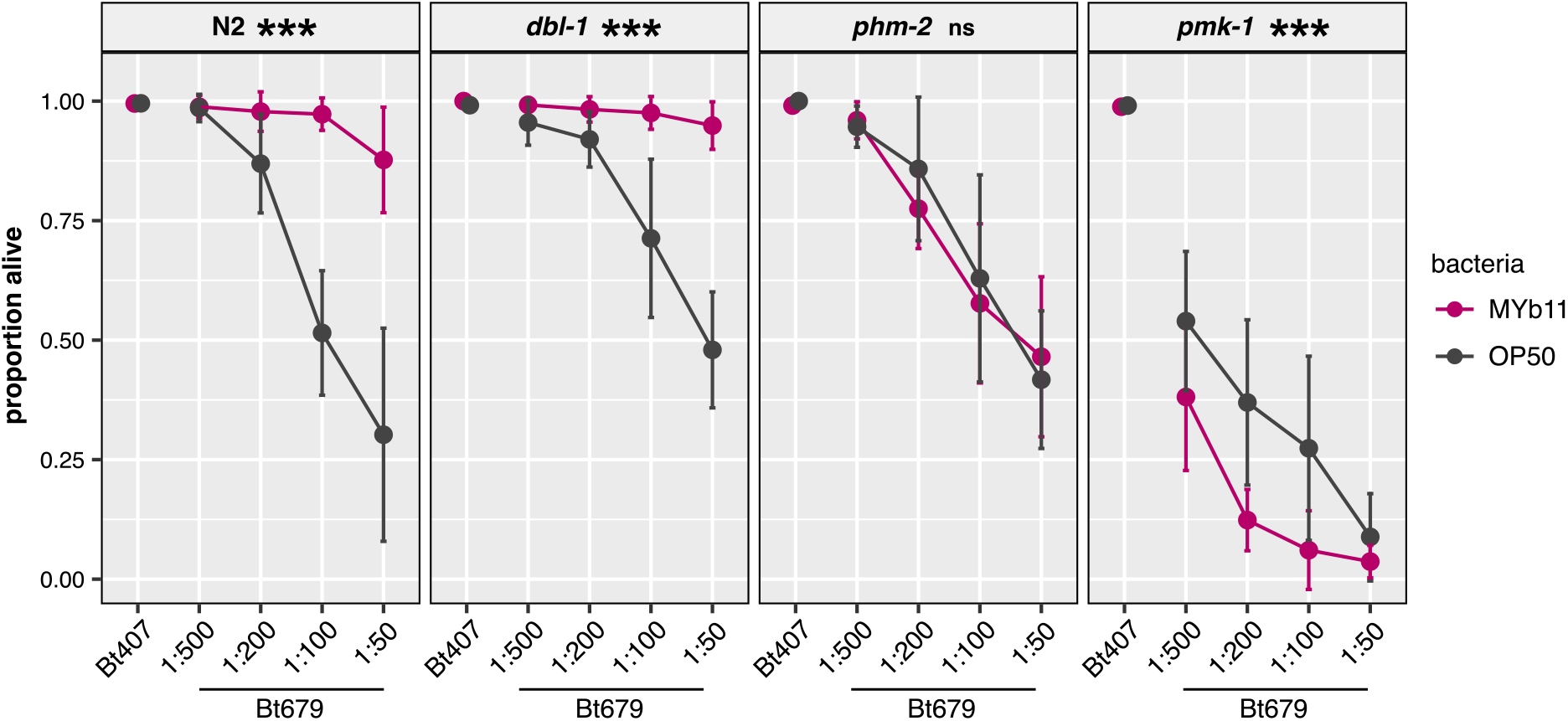
*P. lurida* MYb11 protects *C. elegans* strains N2 and *dbl-1* against infection with Bt679, increases susceptibility of *pmk-1* and has no effect on survival of *phm-2*. Proportion of alive *C. elegans* N2 and *dbl-1, phm-2, pmk-1* mutants fed with MYb11 (pink) or *E. coli* OP50 (grey) 24 h after Bt679 infection. Non-pathogenic Bt407 (1:50) was used as control. Shown are the means as dots and the standard deviations as error bars, with each line representing the proportion of alive worms (survival) on different concentrations of Bt spores. Asterisks denote significant differences between worms on MYb11 and OP50. Generalized linear model, corrected for multiple comparisons with FDR, \*\*\**p* < 0.001, n = 8.

### Interactions between host genetics, protective microbe and pathogen determine *C. elegans* population growth and competitive fitness

We subsequently measured nematode population growth in the presence or absence of Bt679 to determine the combined influence of host genetics, protective microbiota and pathogen as a proxy for metaorganism fitness. All *C. elegans* strains showed decreased population growth on Bt679 compared to non-pathogenic Bt407 (*p* < 0.01) (Figure 3, Supplementary Table S2). Except for *pmk-1* mutants on Bt679, population growth rates were higher on MYb11 compared to on OP50 (*p* < 0.01). On Bt679, the population growth rate of the *pmk-1* mutant was lower than the growth rates of N2 on MYb11 and OP50 (*p* < 0.01). The mutant strains *dbl-1* and *phm-2* always produced lower offspring numbers than N2 on both Bt679 and Bt407 (*p* < 0.05).

**Figure 3:**
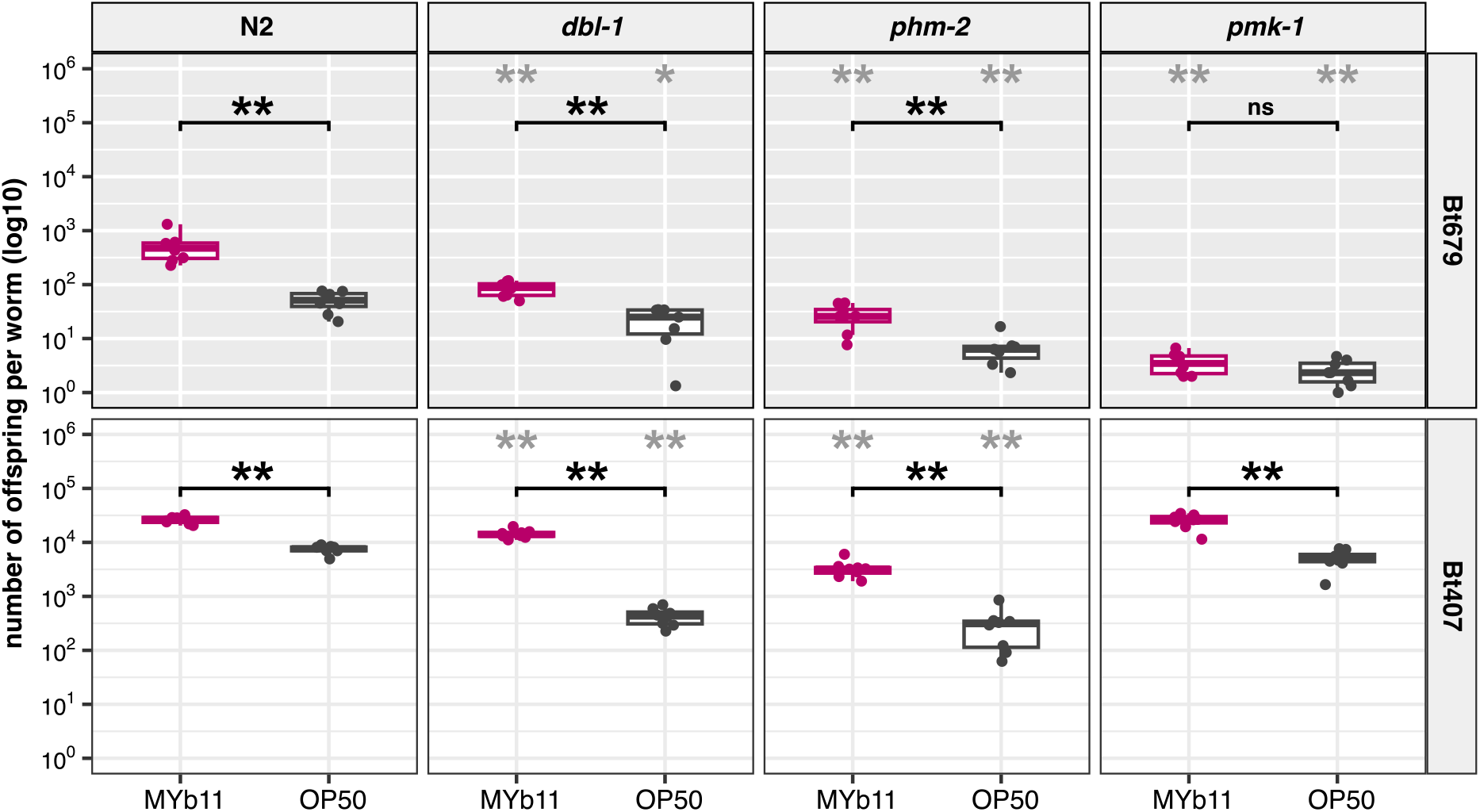
Population growth of *C. elegans* is influenced by host genotype, microbiota and pathogen. The population growth rate of the wildtype N2 and the mutants *dbl-1, phm-2, pmk-1* grown on either *P. lurida* MYb11 (pink) or *E. coli* OP50 (grey) was analysed on pathogenic Bt679 (grey panel background) and non-pathogenic Bt407 (white panel background) and is shown as number of offspring per initial worm. Shown are boxplots with the median as a thick horizontal line, the interquartile range as box, the whiskers as vertical lines, and each replicate depicted by a dot. Asterisks denote significant differences between worms on MYb11 and OP50 (black) or between *C. elegans* mutant strains and wildtype N2 (grey). Wilcoxon Rank-Sum test, corrected for multiple comparisons with FDR, \**p* < 0.05, \*\**p* < 0.01, n = 8.

We further tested the competitive fitness of the mutant strains compared to the GFP-labelled reference strain CL2122 in population growth assays (Figure 4A). The competitive fitness of *C. elegans* CL2122 and N2 did not differ, indicating that the GFP-label does not affect worm fitness (Supplementary Table S1, Supplementary Table S2). The proportions of the *dbl-1* mutant also showed no difference to the reference strain (Figure 4B). *phm-2* mutants on MYb11 and OP50 without Bt679 showed lower competitive fitness compared to the reference strain (*p* < 0.01), and a trend towards lower fitness on OP50 with Bt679 (*p* = 0.082). Except for *pmk-1* mutants, competitive fitness of *C. elegans* on MYb11 and OP50 did not differ. Competitive fitness of *pmk-1* mutants was reduced compared to the reference strain for worms grown on OP50 (*p* = 0.019) and MYb11 (*p* = 0.002) with Bt679, with MYb11 producing even lower fitness than OP50 (*p* = 0.019). Conversely, when Bt679 was absent, then *pmk-1* mutants showed increased competetive fitness on MYb11 compared to on OP50 (*p* = 0.001), and on MYb11, the *pmk-1* mutant produced more offspring than the reference strain (*p* < 0.001).

**Figure 4:**
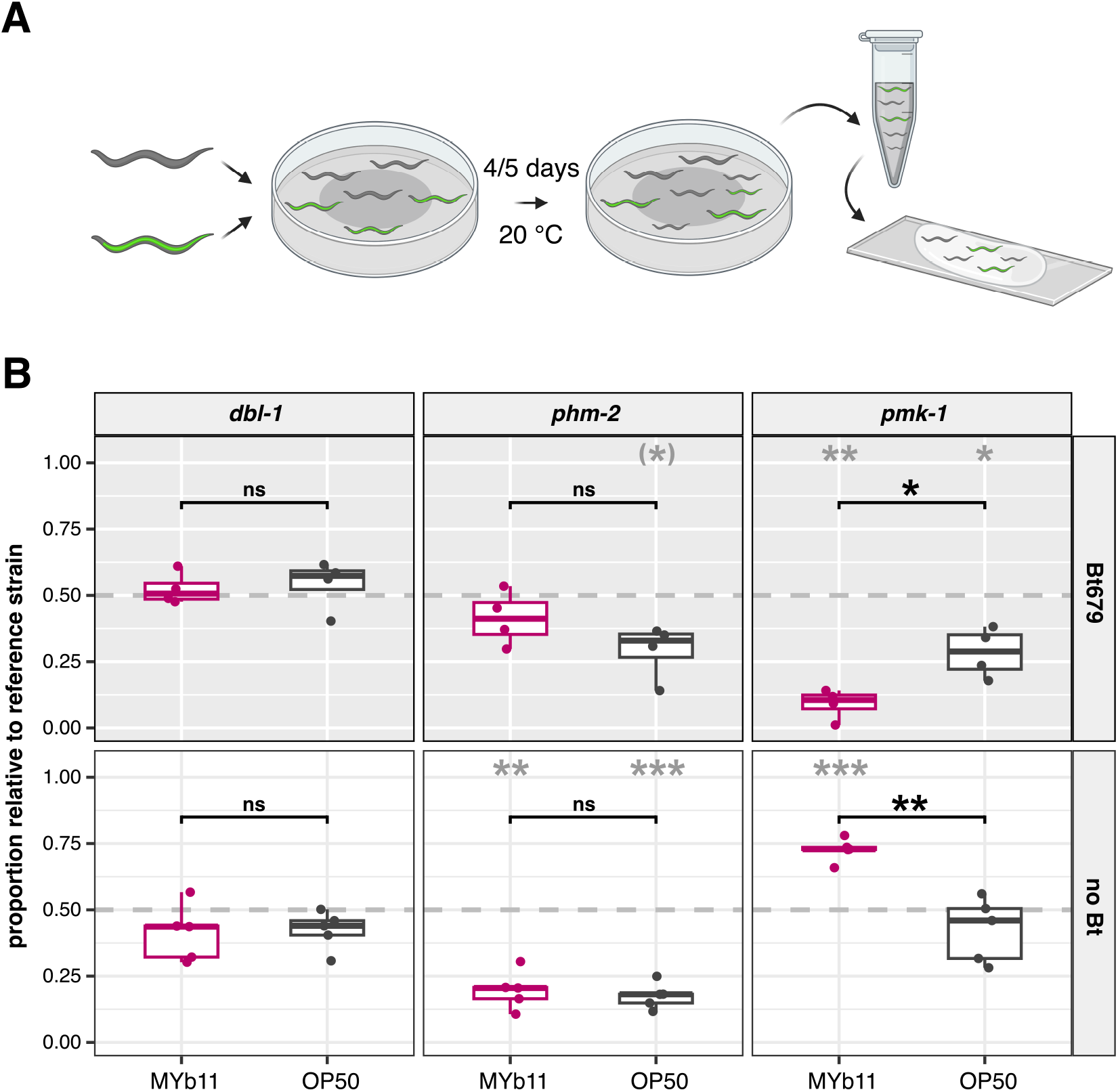
Competitive fitness of *C. elegans* is influenced by host genotype, protective microbe and pathogen. **(A)** Three L4 each of the GFP-labelled reference strain CL2122 and a mutant strain were placed on *P. lurida* MYb11 or *E. coli* OP50 with or without Bt679. After 4 (without Bt679) or 5 days (with Bt679), the relative proportion of mutant worms to the reference strain was determined. **(B)** Proportion of the *C. elegans* mutants *dbl-1, phm-2* and *pmk-1* relative to CL2122 on MYb11 (pink) or OP50 (grey) either with Bt679 (n = 4, with 3 technical replicates each, grey panel background) or without Bt679 (n = 5, with 3 technical replicates each, white panel background). Data are presented as boxplots with the median as a thick horizontal line, the interquartile range as box, the whiskers as vertical lines, and each replicate depicted by a dot. Grey asterisks denote significant differences between the proportion of the respective mutant strain and 0.5 (which is the expected proportion if mutant and wildtype would be present in equal numbers). One Sample t-test, corrected for multiple comparisons with FDR. Black asterisks denote the difference between worms fed with MYb11 or OP50. Two Sample t-test, corrected for multiple comparisons with FDR, ^(^*^)^*p < 0*.*1*, \**p* < 0.05, \*\**p* < 0.01, \*\*\**p* < 0.001.

## Discussion

Our study assessed the triangular interaction between host genetics, protective microbe, and pathogen and their influence on metaorganism fitness. We asked how *C. elegans* mutants interact with the immune-protective microbiota member *P. lurida* MYb11 to determine resistance, population growth and competitive fitness upon infection with the pathogen *B. thuringiensis* Bt679. We used *C. elegans* mutants that affect immunity and/or microbial colonisation in distinct ways, including strains with high colonisation and high pathogen survival (*dbl-1*), high colonisation and low survival (*phm-2*), low colonisation and low survival (*pmk-1*), and also low colonisation and high survival in the wildtype N2. Our main findings are that (i) under pathogen stress, the immunocompetent wildtype N2 generally produces the highest competitive fitness and the highest population growth rate if compared to the mutant strains, suggesting that immunocompetence is more important than high colonisation rates with the protective microbe (as documented for the *dbl-1* and *phm-2* mutants), (ii) association with the protective microbe *P. lurida* MYb11 rather than the standard laboratory food *E. coli* OP50 generally increases host performance under both pathogen and control conditions, suggesting a significant nutritional and/or protective impact of MYb11 on the host, and (iii) disruption of a Bt679-relevant component of the host immune system, the p38 MAPK signalling pathway, completely reverses the effect of the protective microbe, which under pathogen conditions further reduces nematode survival and competitive fitness. In the following, we discuss how the results for each considered mutant enhances our understanding of the worm’s interaction with protective symbionts and pathogens.

The *dbl-1* gene encodes one of four TGF-β ligands in *C. elegans* and it is central for the nematode TGF-β signalling cascade, which is involved in immune defence against fungi and some pathogenic Proteobacteria [67,68]. To date, it was not known to contribute to defence against *B. thuringiensis*. Thus, our observation of an increased survival of the DBL-1 defective mutant on Bt679, without MYb11 and relative to wildtype N2 (Figure 2), suggests a previously unknown role of this pathway in immunity that appears to rely on its inactivation. Moreover, disruption of this pathway causes increased colonisation by MYb11 (irrespective of pathogen environment; Figure 1B). This result indicates that intact TGF-β signalling controls microbiota abundance, as previously reported for other microbiota members in *C. elegans*, including those from the families *Enterobacteriaceae* and *Pseudomonadaceae* [28]. The colonization with MYb11 then leads to increases in survival in the presence of the pathogen (Figure 2) and population growth (irrespective of pathogen environment, Figure 3), suggesting an immune-protective and also a nutritional effect of the symbiont in this genetic background, similarly to what we saw for the N2 wildtype background. Surprisingly, the significantly higher MYb11-colonization rate in the *dbl-1* mutant does not translate into a higher competitive fitness relative to N2 (Figure 4B), indicating that the higher MYb11 colonisation rate alone is not the most critical determinant of nematode fitness.

The results for the *phm-2* mutant confirm that high colonisation rates with the protective symbiont MYb11 do not necessarily improve performance of *C. elegans* in a pathogen environment. The *phm-2* mutant has a defective grinder in its pharynx, which was shown to increase accumulation of alive bacteria in the intestine [69,70]. We similarly observed a significant increase in colonisation by MYb11 and a moderate although insignificant higher abundance of pathogenic Bt679 (Figure 1). The increased MYb11 colonisation did not lead to any change in survival in the presence of the pathogen (Figure 2), but it increased population growth (Figure 3) and also slightly albeit insignificantly competitive fitness (Figure 4), always in comparison to the Bt407 or *E. coli* OP50 control. Population growth and competitive fitness of the MYb11 colonised mutants with the pathogen are either lower or not significantly different from the wildtype N2 measured under the same conditions. Unexpectedly, the competitive fitness of the *phm-2* mutant under pathogen-free conditions is significantly lower compared to wildtype N2. Overall, these results suggest that the defective grinder mutant is generally not as fit as the wildtype, possibly because of less efficient processing of bacterial food. At the same time, it does not suffer significantly more from pathogen infection compared to N2, while the increased colonisation with the protective microbe only enhances offspring production but not survival, thereby revealing a trait-specific effect of the protective microbe in this genotypic host background that differs from the more consistent MYb11-dependent effects in the *dbl-1* mutant background.

The most unexpected results were obtained for the *pmk-1* mutant. The *pmk-1* gene encodes the p38 MAPK and thus a central component of the p38 MAPK signalling cascade, which is one of the main *C. elegans* immunity pathways involved in the defence against diverse pathogen taxa and types [43], including the here used *B. thuringiensis* strain Bt679 [42]. In detail, *B. thuringiensis* produces crystal pore-forming toxins that damage the epithelial membrane of their targets [42,71]. The p38 MAPK pathway plays an essential role in host defence against these toxins [42,72,73] as shown for the Bt toxins Cry5B and Cry21A [39]. Cry21Aa3, a subfamily of Cry21A, is expressed by Bt679 [42,71]. Disruption of the p38 MAPK pathway in *pmk-1* mutants may therefore lead to increased toxin concentrations and epithelial damage, which should cause a decrease in survival, population growth and competitiveness, as observed by us in the current study (Figures 2-4). Intriguingly, MYb11 colonisation was higher in Bt679-infected *pmk-1* mutants, but this effect led to a significant reduction in survival and competitiveness upon pathogen infection. Thus, the protective MYb11 makes it worse for an immunocompromised host in the presence of the pathogen. A possible explanation for this result is the increased damage caused by Bt679 infection in the immunocompromised host, which can promote the translocation of intestinal microbiota into the body cavity, where the microbes may feed on damaged host tissue, enhancing bacterial proliferation, ultimately causing host death, as previously reported for insects infected with insecticidal *B. thuringiensis* [74,75]. MYb11 may thus exploit changes in the host environment to increase its own fitness. These findings support previous work showing that MYb11 need not always be beneficial, but that it has pathogenic potential depending on the context [48]. Moreover, the results highlight the importance of an intact immune response targeted at the infecting pathogen, as provided by the non-disrupted p38 MAPK pathway in the N2 wildtype, as a basis for an additional immune-protective effect of the microbiota, including MYb11, then leading to the better performance of the MYb11-colonised N2 wildtype upon pathogen infection when compared to the corresponding treatments of the *pmk-1* mutant for all measured traits (Figures 2-4). This result is generally consistent with the previous report for *C. elegans*, where expression of a host lysozyme disproportionately suppressed the pathogenic *S. aureus* resulting in enhanced protection by *E. faecalis* [38].

We were surprised to see that the observed population growth rates, measured for each host strain alone, do not directly translate into competitive fitness. More specifically, all mutants produced significantly lower population growth rates than the wildtype N2 under almost all of the tested conditions (Figure 3). This could have been expected for disruption of the *dbl-1* and *phm-2* genes, which was previously reported in both cases to cause reduced offspring numbers [70,76]. The here observed lower population growth rates only led to lower competitiveness of the mutants in four cases, the *phm-2* mutant assayed under pathogen-free conditions and the *pmk-1* mutant under pathogen conditions (always in comparison to N2; Figure 4). This was not the case for the other relevant cases, most notably for the *dbl-1* mutant, which did not differ significantly in competitiveness from N2 under all tested conditions (Figure 4). These results highlight the importance of measuring fitness of a particular genotype not only when it is studied alone but also in competition with other genotypes.

In summary, our study demonstrates that the host genotype critically determines the beneficial effects of a protective symbiont in the presence of a pathogen. In certain genomic contexts, the symbiont can provide protection, whereas in other genomic contexts it can further decrease survival and fitness, which we observed in mutants with a disrupted immune response towards the considered pathogen. Therefore, the particular characteristics of the triangular interaction between host genes, protective microbes and pathogen need to be considered for a full understanding of metaorganism fitness. The here described relationships have been inferred with a simplified model and, thus, they are likely even more complex under realistic conditions with a much more speciose and dynamic microbiota community.

## Acknowledgements

We thank the Schulenburg group for helpful feedback, and the Caenorhabditis Genetics Center (CGC), funded by the National Institutes of Health (NIH) Office of Research Infrastructure Programs (P40 OD010440), for providing the *C. elegans* strains and *E. coli* OP50. Illustrations for experimental designs were created using BioRender.com.

## Data Availability Statement

All data generated or analysed during this study are included in this published article and its supplementary information files.

## Author contributions

HGK: Methodology, Investigation, Formal analysis, Writing-original draft. CP: Methodology, Investigation, Formal analysis, Writing-original draft. IKH: Methodology, Investigation, Writing-review & editing. HS: Conceptualisation, Writing original draft, Funding acquisition.

## Supplementary Figure

**Supplementary Figure S1:**
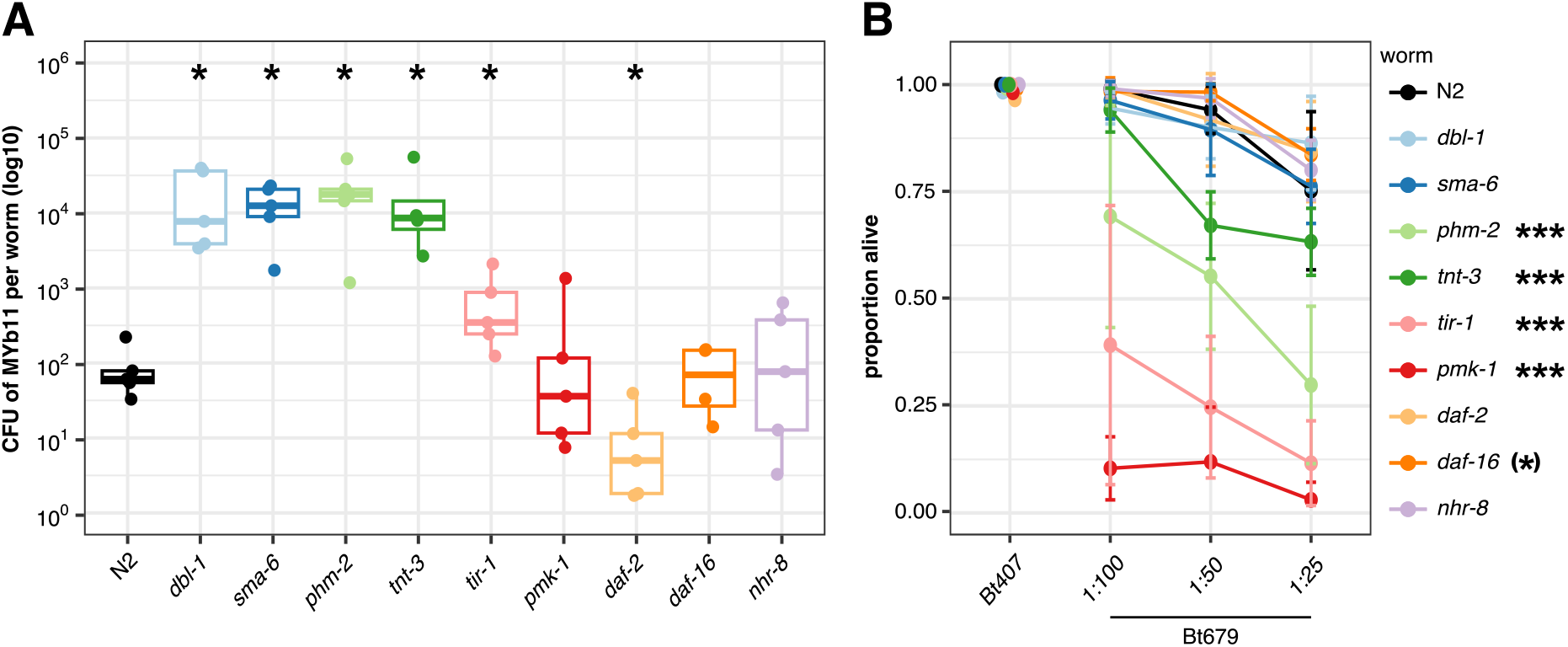
Individual *C. elegans* genes influence colonisation rates with *P. lurida* MYb11 and survival on *B. thuringiensis* Bt679. **(A)** MYb11 colonisation rates in one-day old adult *C. elegans* shown as colony forming units (CFU) per worm in boxplots with the median as a thick horizontal line, the interquartile range as box, the whiskers as vertical lines, and each replicate depicted by a dot. Note that the y-axis is log10-transformed. Asterisks denote differences between *C. elegans* mutant strains and the wildtype N2. Wilcoxon Rank-Sum test, corrected for multiple comparisons with FDR, \**p* < 0.05, n = 5. **(B)** Survival of *C. elegans* fed with MYb11 24 h after infection with Bt679. Non-pathogenic Bt407 (1:25) was used as control. Shown are the means as dots and the standard deviations as error bars, with each line representing the proportion of alive worms (survival) on different concentrations of Bt spores. Asterisks denote differences between *C. elegans* mutant strains and the wildtype N2. Generalized linear model, corrected for multiple comparisons with FDR, ^(^*^)^*p* < 0.1, \*\*\**p* < 0.001, n = 4.

