## Supplementary material for "The intricate triangular interaction between protective microbe, pathogen, and host genetics determines fitness of the metaorganism": Supplemantary Information

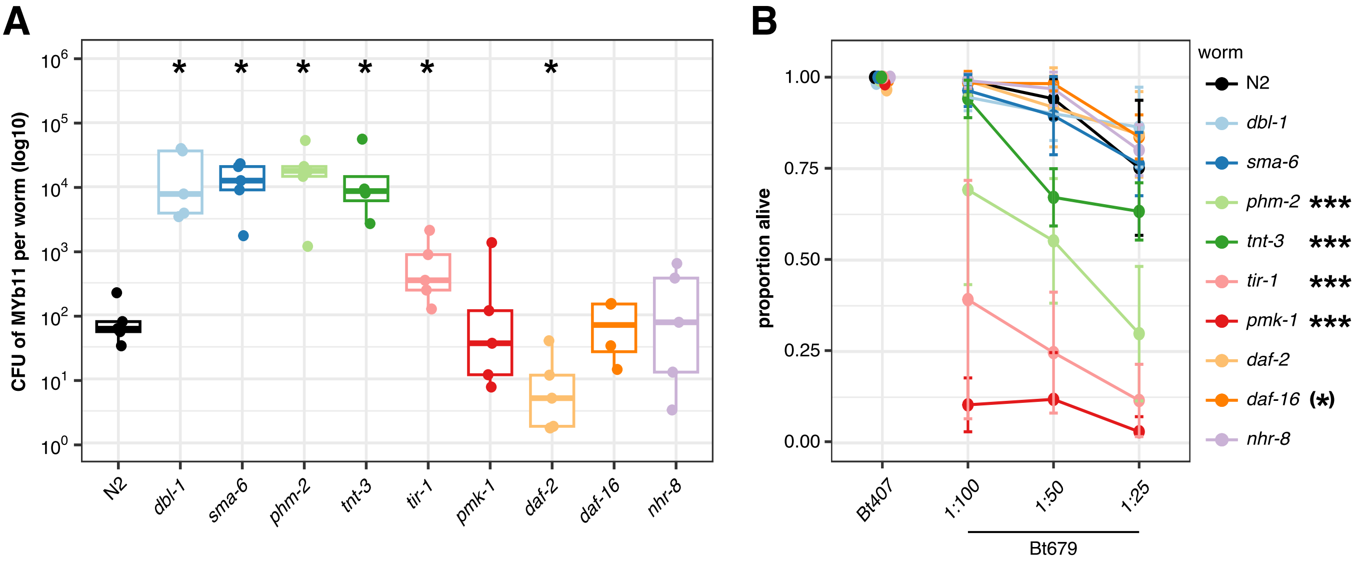


**Supplementary Figure S1: Individual *C. elegans* genes influence colonisation rates with *P. lurida* MYb11 and survival on *B. thuringiensis* Bt679. (A)** MYb11 colonisation rates in one-day old adult *C. elegans* shown as colony forming units (CFU) per worm in boxplots with the median as a thick horizontal line, the interquartile range as box, the whiskers as vertical lines, and each replicate depicted by a dot. Note that the y-axis is log10-transformed. Asterisks denote differences between *C. elegans* mutant strains and the wildtype N2. Wilcoxon Rank-Sum test, corrected for multiple comparisons with FDR, $^{*}p < 0.05$, n = 5. **(B)** Survival of *C. elegans* fed with MYb11 24 h after infection with Bt679. Non-pathogenic Bt407 (1:25) was used as control. Shown are the means as dots and the standard deviations as error bars, with each line representing the proportion of alive worms (survival) on different concentrations of Bt spores. Asterisks denote differences between *C. elegans* mutant strains and the wildtype N2. Generalized linear model, corrected for multiple comparisons with FDR, $^{(*)}p < 0.1$, $^{***}p < 0.001$, n = 4.
